# Uncovering the genetic blueprint of the *C. elegans* nervous system

**DOI:** 10.1101/2020.05.04.076315

**Authors:** István A. Kovács, Dániel L. Barabási, Albert-László Barabási

**Affiliations:** Department of Physics and Astronomy, Northwestern University, Evanston, IL, USA; Network Science Institute and Department of Physics, Northeastern University, Boston, MA, USA; Wigner Research Centre for Physics, Institute for Solid State Physics and Optics, Budapest, Hungary; Department of Network and Data Science, Central European University, Budapest, Hungary; Biophysics Program, Harvard University, Cambridge, MA, USA; Division of Network Medicine and Department of Medicine, Brigham and Women’s Hospital, Harvard Medical School, Boston, MA, USA

## Abstract

Despite rapid advances in connectome mapping and neuronal genetics, we lack theoretical and computational tools to unveil, in an experimentally testable fashion, the genetic mechanisms that govern neuronal wiring. Here we introduce a computational framework to link the adjacency matrix of a connectome to the expression patterns of its neurons, helping us uncover a set of genetic rules that govern the interactions between adjacent neurons. The method incorporates the biological realities of the system, accounting for noise from data collection limitations, as well as spatial restrictions. The resulting methodology allows us to infer a network of 19 innexin interactions that govern the formation of gap junctions in *C. elegans*, five of which are already supported by experimental data. As advances in single-cell gene expression profiling increase the accuracy and the coverage of the data, the developed framework will allow researchers to systematically infer experimentally testable connection rules, offering mechanistic predictions for synapse and gap junction formation.

## Introduction

There is ample experimental evidence that the connectome, capturing the neuron level wiring of a brain, is genetically encoded. Indeed, while neurons are clustered into broad classes based on their morphology and function, these observed differences between cells are known to be rooted in the differential expression patterns of their genes and proteins [1–9]. Consequently, perturbations that alter the genetic identity of individual neurons can induce significant changes in wiring [10, 11]. Furthermore, developmental neuroscience has unveiled multiple genetic factors contributing to the formation of neuronal circuits. For example, the connectome of *C. elegans* and higher organisms rely on a combination of body and wiring localization [12–17], and cell-cell recognition specificity, both for synaptic [18] and gap junction connections [10, 19, 20]. In the mouse retina, proteins, like connexin-36, play a known role in coupling rods and cones through gap junctions [21] and in *D. melanogaster*, neurons expressing the same olfactory receptor converge onto the same set of projection neurons [22]. While these studies offer strong experimental support for the genetic roots of neuronal wiring, we continue to lack a general framework to identify the genetic mechanisms that determine the presence or the absence of specific neuronal connections [11, 20, 23].

These advances have prompted the development of statistical approaches designed to identify genes involved in synaptic connectivity [24–26]. Notably, Kaufman et al. [24] demonstrated a correlation between gene expression and neuronal connectivity, and Varadan et al. [25] identified a genetic rule for chemical synapses through an entropy minimization approach. However, these frameworks do not incorporate the constraint that synapses can only exist between physically adjacent neurons, prompting Baruch et al. [26] to estimate spatial proximity information based on neuron connectivity pattern. Notwithstanding these promising advances, progress towards unveiling the genetic rules of synapse formation is remarkably slow compared to the tremendous experimental progress focusing on mapping the connectome and the gene expression patterns of individual neurons [27–29].

The gap between experimental and computational progress raises a fundamental question: is it computationally feasible to infer the genetic rules that govern synapse formation from the available experimental data? For instance, in *C. elegans* we wish to describe the genetic rules that govern the wiring of neurons of a relatively sparse connectome of *N* ∼ 300 neurons [27] using as input the combinatorial expression patterns of *m* ∼ 20, 000 genes [29]. In general, if in each neuron m genes contribute to synapse formation, together they can describe a connectome of up to *N* = 2^*m*^ neurons. Hence, as we try to infer the gene list whose expression pattern can explain the observed connectome, we are faced with a heavily under-determined problem: in *C. elegans* a randomly generated expression of *m* = log2(*N*) < 9 genes can fully describe the observed connectome without revealing any biological information. Although humans have *N* ∼ 86 billion neurons [30], and only *m* ∼ 20, 000 genes, the number of neurons is dwarfed by the combinatorial gene expression space of size 2^*m*^, where the expression pattern of *m* genes determines whether two neurons can synapse. Indeed, if only three genes contribute to synapse formation in each neuron in the human brain, they allow for 1/6 × *m*^3^ = 1.3 × 10^12^ combinations, an order of magnitude larger than the number of neurons in a human brain, leading again to serious overfitting. We are therefore faced with an astronomical search space, and the challenge to extract meaningful genetic rules in a heavily ill conditioned problem of finding them from inherently limited experimental data.

In this paper we show how to overcome these difficulties, relying on network and physical approaches that are known to provide complex structures from simple processes [31–35]. We begin by developing a theoretical and modeling framework to systematically infer the genetic rules that contribute to the formation and maintenance of synapses and gap junctions between adjacent neurons. We then show that these genetic rules can be systematically extracted from two datasets: (i) a comprehensive map of the connectome and (ii) a protein expression atlas of the individual neurons. Finally we rely on the roundworm *Caenorhabditis elegans* to test our modeling framework. We do so because the *C. elegans* connectome is believed to be largely identical across individuals [28, 36, 37], hence fully predetermined by the genetic markers that label each neuron [19, 38]. Yet, the genetic mechanisms that determine which neurons can synapse with each other remain largely unknown even in this simple and well-studied organism [20].We show that we can overcome overfitting by restricting our analysis to genes known to be involved in gap junction formation. We demonstrate the utility of the proposed modeling framework by predicting 19 interactions between innexin proteins responsible for gap junction formation, finding that 5 of them are supported by previous experimental data. Finally, we show that the SCM reveals the non-Euclidean organization of brain wiring, indicating that the *C. elegans* connectome is not driven, but is merely constrained by spatial factors.

### The Connectome Model

We begin with two hypotheses, the first being that each gene can be in two possible states, expressed (1) or not (0), whose combination define the genetic barcode for each neuron. As synapses form between pairs of neurons, the second hypothesis states that synapse formation is governed by some unknown biological mechanism linked to the gene expression pattern of each neuron (neuronal barcodes). We describe each such a mechanism as an operator *O*, which inspects the barcodes of two neurons and decides to facilitate (or block) the formation of synapses or gap junctions between them [39].

Consider a hypothetical connectome consisting of seven neurons, A-G, whose connections are uniquely determined by the expression patterns of three genes (Fig. 1a). The Connectome Model consists of a set of rules that encode the possibility of synapses between genetically encoded sets of neurons [39]. In the simplest case, a rule could be an operator *O*_1_ that recognizes the complete genetic profile of neurons C and G, designating C as a source and G as a destination neuron, and establishing synapses between them (Fig. 1b). However, a less specific operator (*O*_2_ or *O*_3_), that detects only a subset of the genes, ignoring the expression state of the genes marked by *X*, can generate multiple links between two sets of neurons, like the complete biclique of eight links in Fig. 1d. Figure 1 summarizes a key prediction of the Connectome Model: Each biological mechanism that relies on gene expression to initiate synapse formation will generate an imprint in the connectome in the form of a unique network motif, known as a *non-induced biclique* in graph theory [40] (see also Supporting Figure S1), where all neurons of the source set can be connected to all neurons of the destination set. Ref. [39] validated this prediction by showing an excess of specific large biclique motifs in the *C. elegans* connectome. The challenge, which we address here, is how to reverse engineer the genetic rules from the observed network patterns, given that even a modest number of genetic rules can lead to a tremendous number of network motifs. Furthermore, the genetic rules can be rather complex when expressed in terms of operators connecting gene expression patterns (Fig. 1d), rooted in the non-linear representation of combinatorial expression data. To reduce this complexity, we introduce genetic *labels*, allowing us to capture multiple genetic operators within a single network description. To be specific, for an operator *O*_*ab*_, we assign all participating source neurons a label (“a” in Fig 1e) and all destination neurons a different label (“b” in Fig. 1e). *A* biological rule governing neural connections between the source and destination neurons can be represented by the link a − b between the two labels. In this label-based representation, the operators have a simple form (Fig. 1e-g).

**FIG. 1:**
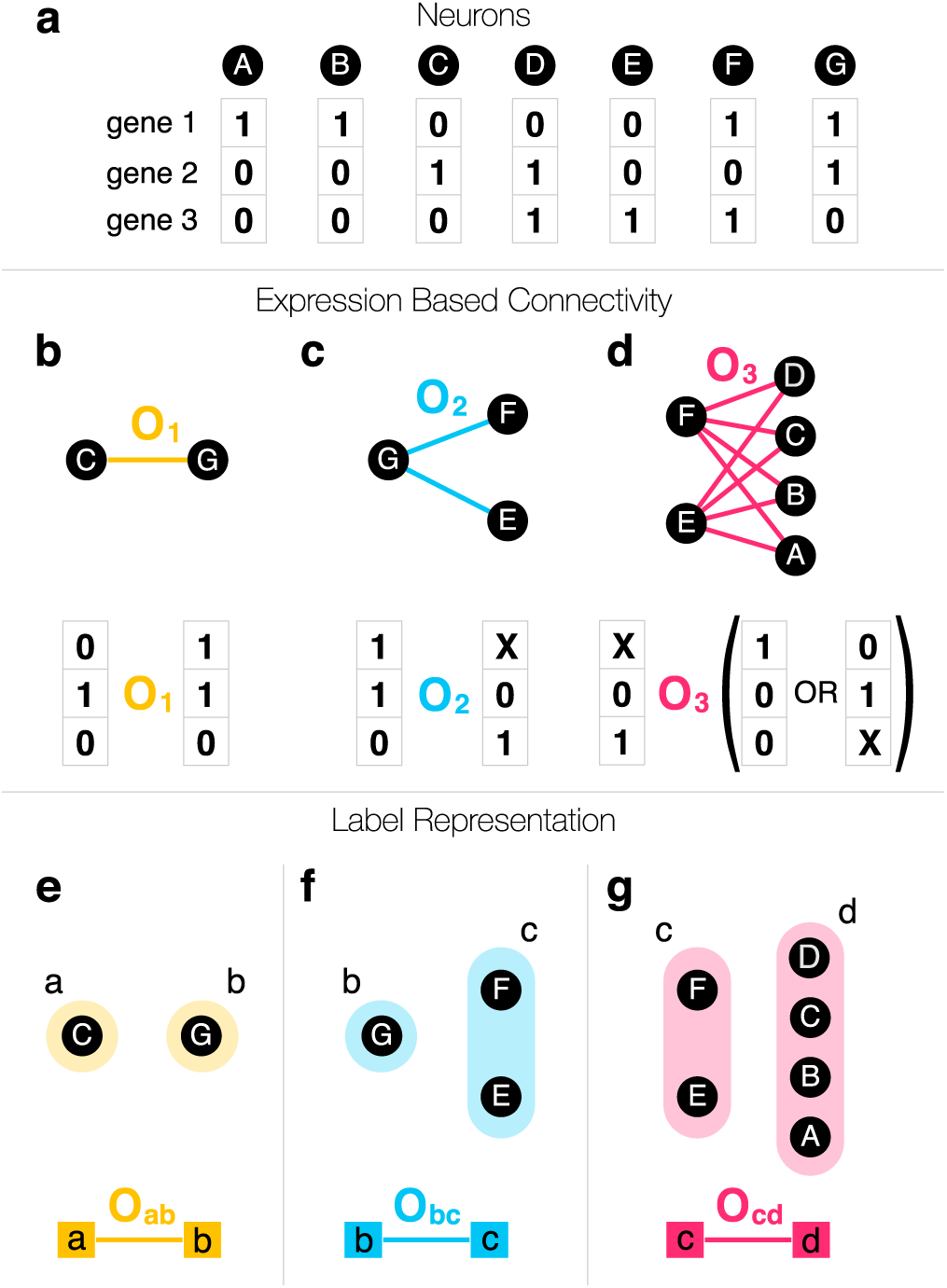
Gene Expression and the Connectome. **a)** As neural connections are determined by neuron identity, we start with the terminal expression profile of seven neurons (blue nodes), whose connections to each other is determined by the expression patterns of three genes, expressed (1) or unexpressed (0). **b)** The formation of links (chemical synapses, gap junctions) are determined by the expression profiles of the neurons, driven by biological mechanisms that are abstracted as operators, *O*_*i*_. In the simplest case, an operator recognizes the full expression pattern of neurons C and G and connects them. **c)** *A* single rule or operator can generate multiple links if the operator is more restricted, i.e. detects the expression of some genes and ignores others. Here, X marks the gene ignored by the operator, whether it is expressed or not. **d)** More specific operators that have multiple X’s in them can facilitate a large number of links, see Eq. (2). **e)** To each operator we assign two labels, one to the source neurons (left) and another to the destination neurons (right). The labels allow us to represent operator *O*_1_ as a link connecting the neurons with the right labels. **f)** Even if the same label is assigned to multiple neurons, the operator *O*_2_ remains a simple link between the two labels. **g)** While the operator *O*_3_ might appear complicated in terms of the original gene expression data, it has a simple structure in the label representation.

Although some labels can represent complex gene expression patterns (e.g. Fig. 1d) others can be very simple. For example, electrical synapses or gap junctions (GJs) are inter-cellular channels formed by two matching hemi-channels consisting of a subset of 25 innexin proteins [41]. In order to maintain a GJ, the innexins forming the hemi-channels must be expressed on the surface of adjacent neurons. In our formalism the simplest rule, *O*_*ab*_, is then a link between two innexins “a” and “b” expressed on two neurons forming a GJ. In other words, we assign label “a” (or “b”) to each neuron if it expresses innexin “a” (or “b”).

The labelling of each neuron is summarized in the expression matrix (*X*), where *X*_*ia*_ = 1 if neuron i’s expression pattern is consistent with label “a”, and zero otherwise (Fig. 2a). The rule matrix *O* summarizes the individual operators as links connecting the labels (Fig. 2b). If two adjacent neurons express labels that are connected in *O*, then there is a non-zero chance of establishing a synapse between these neurons. This representation defines mathematically the Connectome Model (CM), that links the brain’s connectome (*B*) to the expression patterns of the individual neurons *X*, through the rule matrix *O*,

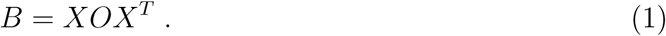

**FIG. 2:**
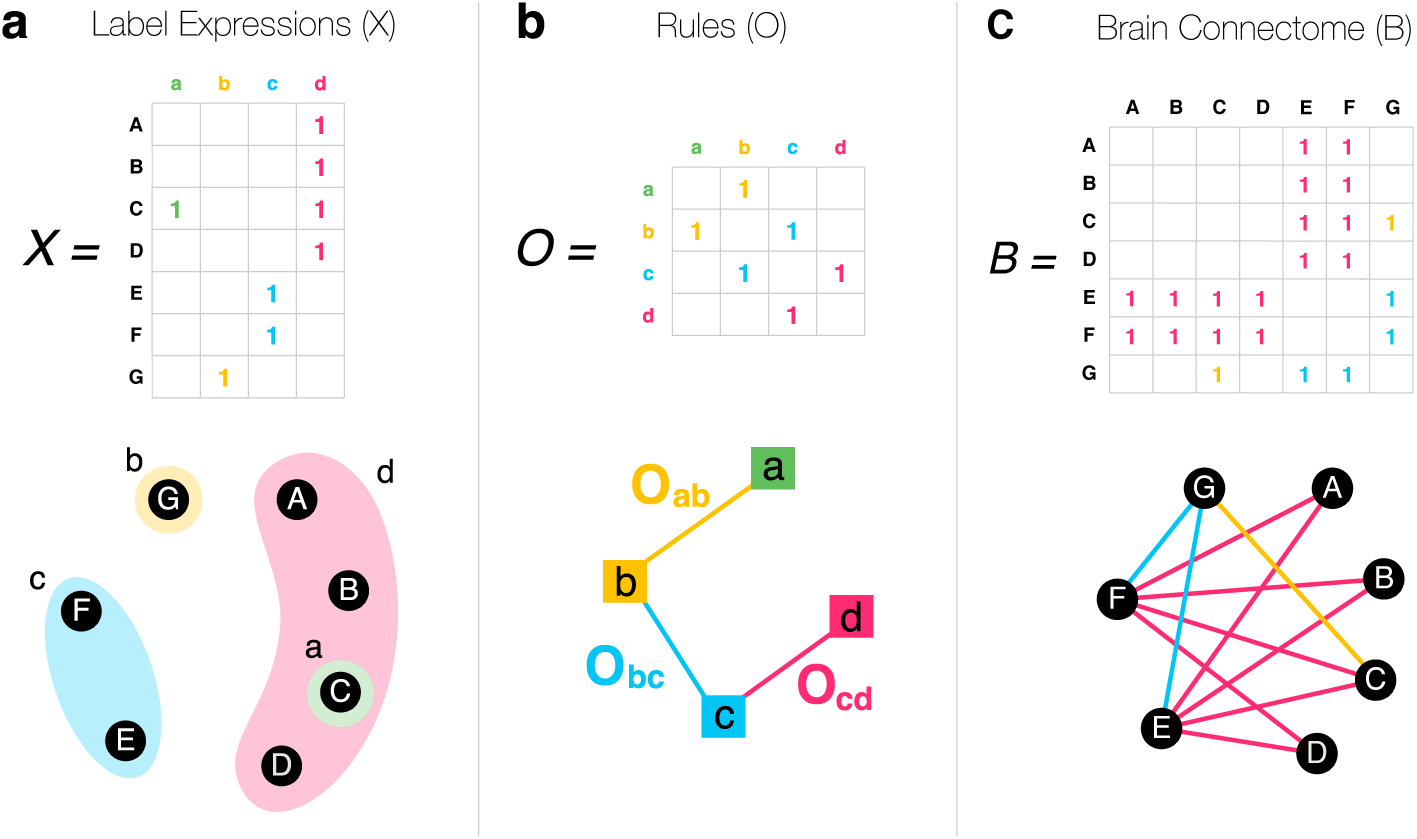
The Connectome Model (CM). **a)** The expression pattern of the neurons A-G are summarized in the label expression matrix *X*. **b)** The operators *O*_1_ − *O*_3_ connecting the labels can be summarized in the organizing rule matrix *O*. **c)** In the CM, the brain connectome (*B*) emerges from *O* and *X* through the CM Eq. (1). Each time two labels are connected in *O*, the corresponding neurons in *X* form synapses. Only non-zero elements are shown in the matrices.

The brain equation (1) is our first key result, formally linking the connectome (*B*), the expression patterns of the individual neurons (*X*), and the biological mechanisms (*O*) that govern synapse/GJ formation in the brain.

In practice, not all of the genetically allowed connections can be observed, due to experimental limitations, developmental and spatial constraints as well as neural plasticity. In our formalism this implies that *O* is a stochastic operator with *O*_*ab*_ < 1 (Fig. 3). For instance, fruit fly inx-2 homomeric GJs form only between 40% of the neighboring cell pairs [42], leading to an apparent *stochasticity* in GJ formation. In the absence of such stochastic effects, *O*_*aa*_ predicts a complete subgraph of all a label neurons. With stochasticity, instead of a fully connected subgraph, we expect a community of nodes connected to each other with density *O*_*aa*_ = 0.4 (Fig. 3e). In other words, *O* is a weighted matrix, where the weights are the probabilities that neurons carrying labels “a” and “b” will link to each other.

**FIG. 3:**
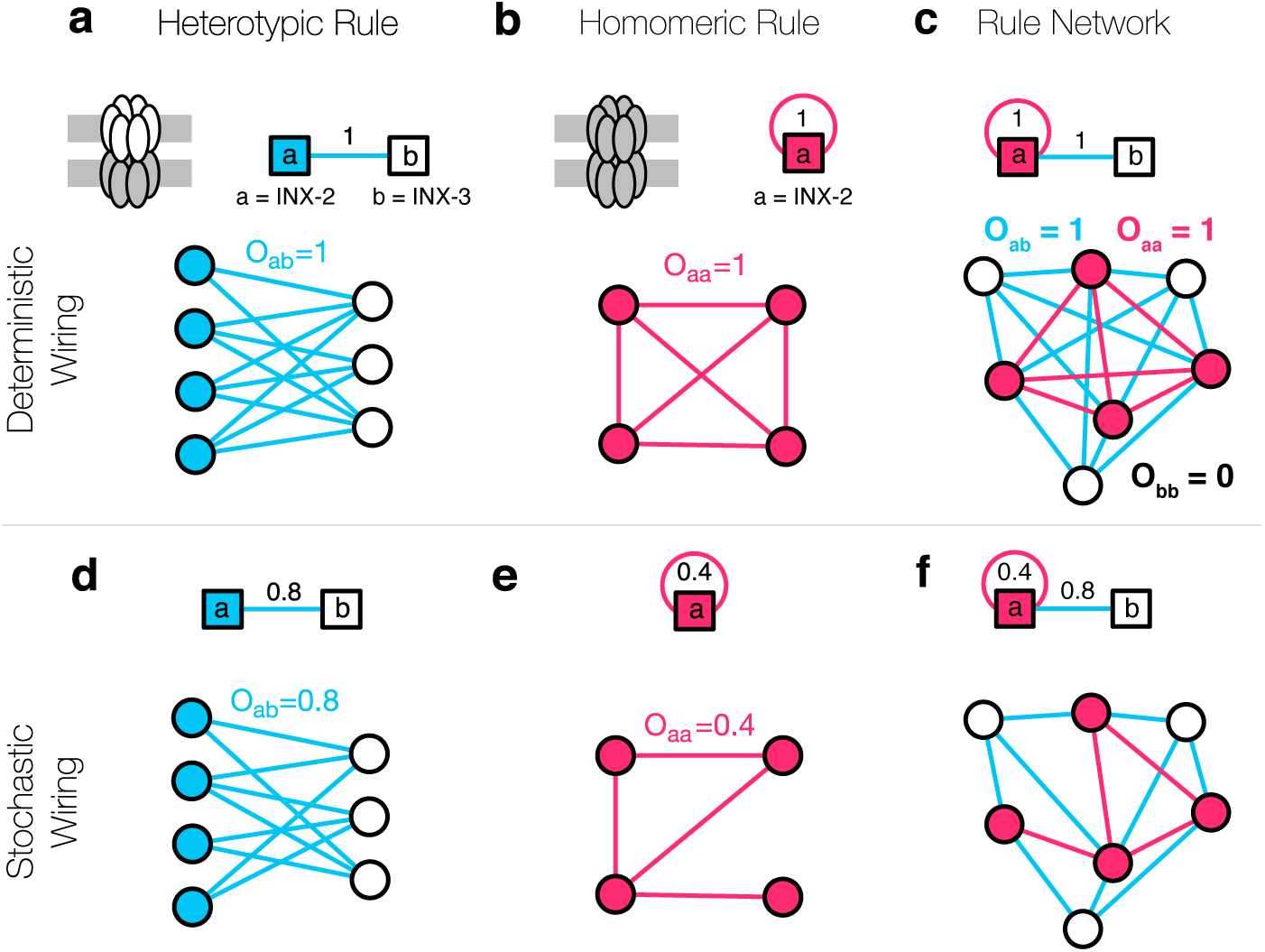
Mapping Gap Junctions to the Connectome Model. Gap-junctions are formed by interacting hemi-channels comprised of innexin proteins. In the simplest case, a hemi-channel is made of a single innexin, meaning that the expressed innexins can directly serve as labels. **a)** Two Drosophila innexin proteins, inx-2 and inx-3 have been found to form (heterotypic) gap-junctions, resulting in multiple potential neural connections [42]. **b)** There is evidence that inx-2 can form homomeric gap-junctions, establishing connections between the neurons expressing inx-2, represented by the self-loop in the figure. **c)** Altogether, the two rules (a) and (b) can be integrated into a rule network that serves as a template for the entire gap-junction connectome. **d)** The formalism behind the CM allows for stochastic rules, i.e. a weight or 0.8 indicates that 80% of the potential neural connection are present in the brain. This stochasticity can arise from multiple factors, including noisy or incomplete expression and connectome data, spatial effects, biological constraints, and the true stochasticity of neuronal wiring. **e)** According to oocyte experiments [51] the homomeric innexin rule of Drosophila inx-2 has a weight of 0.4, as only 40% of the possible links are observed. **f)** Even in the presence of apparent or true stochasticity, we can capture the entire GJ connectome using only a few innexin rules.

Taken together, as the brain equation (1) establishes a direct connection between the expression profiles of the individual neurons (*X*) and the connectome (*B*) through genetic rules (*O*), it allows us to address several key problems in brain science, listed in the order of increasing technical difficulty:

I. *Map out the Connectome:* Predict the connectome (*B*) from gene expression (*X*) and the genetic rules (*O*);
II. *Unveil the Genetic Rules:* Predict the genetic rules (*O*) behind the connectome from known *X* and *B*;
III. *Predict Expression Patterns:* Find the gene expression of neurons (*X*) from the genetic rules (*O*) and the wiring of the connectome (*B*);

Problem I is readily solved by Eq. (1), assuming that we know (some of) the biological mechanisms behind the rules in *O*. As we currently lack these rules, in this paper we focus on the pressing issue of solving Problem II. This choice is motivated by the fact that in *C. elegans* we have a comprehensive map of its neural system’s adjacency matrix (*B*) and extensive (yet somewhat noisy and incomplete) information on the gene expression patterns of individual neurons (*X*), potentially allowing us to determine the biological mechanisms ecnoded in *O*.

### Solving the Connectome Model

Given the connectome *B* and the labels *X*, our goal is to identify the operator *O* that collects the biological rules that govern link formation (Problem II). To illustrate the procedure, we use the three rules introduced in Fig. 1 to generate the brain connectome *B* (Fig. 2c), according to the label expression *X* (Fig. 2a). Just by looking at two connected neurons it appears impossible to reverse the problem and infer the genetic rule responsible for each connection (Fig. 2a, neurons C and G). Indeed, the rule could connect label “a” to label “b”, but could also connect label “a” to label “d”. Even if we had simultaneous access to the complete list of neural connections (*B*) and full genetic labels (*X*), inferring the rules responsible for link formation (*O*) is mathematically ill conditioned, with infinitely many solutions of the form

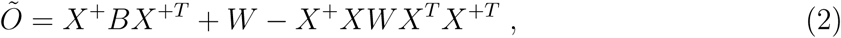

where *W* is an arbitrary matrix and *X*^+^ stands for the Moore-Penrose pseudo-inverse of *X*. We have a unique solution only when *X*^+^ = *X*^−1^, meaning that the neurons have linearly independent expression patterns, which is not expected to be the case in the brain. Otherwise, even if there is no noise in the input data, we do not expect to find exact solutions, and *Õ* comes with a least-square residual error *r*^2^ = ||*B* − *XÕX*^*T*^ ||^2^ > 0. In practice, the situation is even more difficult because *B* and *X* have multiple unknown errors (both false negatives and false positives). To make progress, we invoke the parsimony principle, searching for the model that accounts for the available data with the fewest rules in *O*, i.e. requires the smallest possible number of distinct biological hypotheses. *A* convenient way to mathematically formalize this is to minimize the objective function with a regularization parameter *α* ≥ 0,

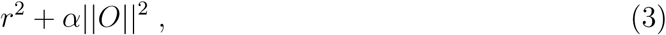

where 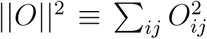 is the square of the Frobenius norm. When *O* consists of only 0s and 1s, a minimal Frobenius norm corresponds to the fewest rules or the fewest 1’s in the *O* matrix. As an alternative implementation of the parsimony principle, we could also select the sum of the absolute values in *O* as the norm, related to compressed sensing, also known as LASSO [43]. Here, we proceed with the Frobenius norm in order to maintain the analytical tractability of the problem, and to be able to assess the significance of the obtained rules. With this, we can find the optimal *O*, relying on the results on ridge regression (Tikhonov regularization) [44], discussed in Methods.

### The Spatial Connectome Model (SCM)

The CM assumes that each neuronal connection allowed by the genetic profile of the neurons will form. Yet, for a synapse or GJ to form, the neurons must also be spatially adjacent (Fig. 4a). By ignoring spatial constraints, each missing link between remote neurons is taken as evidence against the rule, including links allowed by the genetics that do not have the opportunity to form as the neurons do not come close to each other (Fig. 4b). Therefore, to increase the accuracy of the model’s predictions, we need to restrict our analysis to pairs of spatially adjacent neurons. This information is encoded by the spatial adjacency matrix *A*, telling us which neuron pairs are in physical proximity. In the *C. elegans* anterior brain *A* has been mapped experimentally [45], hence we restrict our analyses to the anterior 185 neurons. Here, 5, 592 pairs are adjacent, representing ≈ 33% of all pairs, out of which only 601 form GJs.

**FIG. 4:**
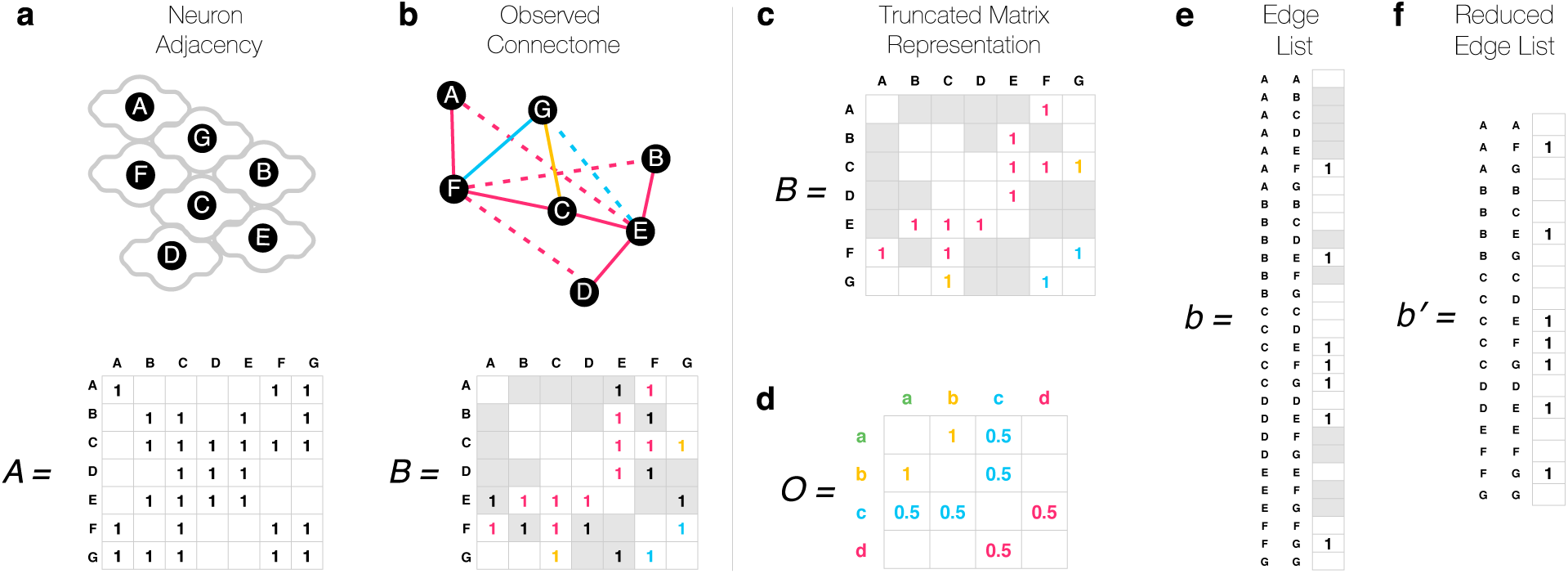
The Spatial Connectome Model. **a)** Neurons can only synapse if they are located in each other’s vicinity. We schematically indicate spatial adjacency via touching neuron contours in the figure, and adjacent neuron pairs are marked by a 1 in the spatial adjacency matrix *A*. **b)** Given the lack of proximity, only a fraction of the genetically allowed synapses are observed. The dashed links in the network, shown as 1s in gray cells in the adjacency matrix below, indicate neural connections that are genetically permitted but are not observed because the neurons are not adjacent. **c)** When inferring genetic rules, distant neuron pairs must be ignored in the model (gray cells), as we do not know if the lack of connection has a genetic origin, or is it simply due to spatial constraints. We therefore arrive to a truncated matrix representation, which do not obey standard matrix operations, and hence are challenging to work with. **d)** If we treat all unobserved cells (gray and blank) as zeros, the matrix representation leads to incorrect rules, as it always assumes the lack of genetic compatibility where there may be some. **e)** The edge list representation offers a linear description that is formally equivalent with the matrix representation. **f)** Distant pairs of neurons can be removed from the edge list representation, and as a truncated list is still a list, it allows us to uncover the correct rules using Eq. (5).

It is tempting to incorporate spatial constraints into our matrix representation (1) by ignoring each matrix element in *B* that is absent in the spatial adjacency matrix *A* (Fig. 4c). If we do so, the obtained truncated matrix has non-existing entries (Fig. 4c), and we cannot apply standard matrix operations to it. Alternatively, treating each missing link as a zero in the connectome matrix leads to incorrect rules, as illustrated in Fig. 4d. To address this problem, we represent the connectome *B* as an edge list rather than a matrix. In other words, we rearrange the connectome matrix *B* (by any, i.e. lexicographic order) into the connectome vector *b* = vec(*B*) (Fig. 4e). Similarly, we rearrange the rule matrix *O* into a rule vector *o* = vec(*O*). This allows us to reformulate the bi-linear CM in Eq. (1) as a higher dimensional linear model

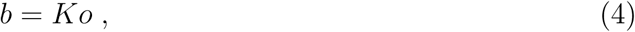

where *K* = *X* ⊗ *X*. In this, so far equivalent, linear representation we can now restrict the space of neural connections to spatially adjacent neurons by ignoring the entries in *b* and *K* that do not satisfy spatial proximity according to the *A* matrix (Fig. 4f). We therefore arrive at the truncated brain equation describing the spatial connectome model (SCM), representing our second key result,

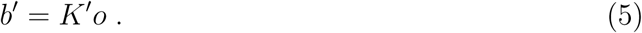

This equation can be solved using tools similar to the ones we used to study the CM, as discussed in Methods. At the end, the obtained rule weights can be rearranged into the matrix format *Õ*′. If we perform these calculations on the toy model of Fig. 2a,c with the indicated spatial constraints, we recover the exact rules in Fig. 2b, even though we are using only a fraction of the connectome information, i.e. only the links that are between adjacent neurons. This result suggests that we do not need complete input data on the *C. elegans* connectome and gene expression to make progress, as we can use Eqs. (5) (and (7) in Methods) to uncover the biological mechanisms *O* governing brain wiring even from partial data. Yet, we need to know the genetic labels, i.e. the genetic basis, X, in which the organizing rules operate. Next we show how (4) helps us unveil the biological mechanism governing gap junction formation.

### Unveiling the Gap Junction Operators

Electrical synapses, or GJs, play an important role in the *C. elegans* nervous system and muscle control [46]. There are 25 genes involved in *C. elegans* GJ formation, all of which encode innexin proteins (collectively called innexin genes, even though not all of them are named inx). We can therefore ignore the expression patterns of non-innexin genes, limiting over-fitting by restricting the genetic space in *X* used in our analysis. Currently, there is published expression data for 18 of the 25 innexin genes in *C. elegans* neurons [45]. Although every neuron class is known to form GJs, about one third of the neurons have no reported innexin gene expressed [47, 48]. Besides this obvious data incompleteness, the expression data is also limited by experimental difficulties of differentiating between individual neurons within the same neuron class, limiting the resolution of the expression data to neuron classes. With these data limitations in mind, as a first step, we consider only the genetic labels linked to the expression patterns of individual innexin genes.

We begin by applying Eq. (5), using as input the innexin gene expression data (*X*) [45], the GJ connectome (*B*) [28] and the spatial adjacency *A* [29, 45], aiming to calculate *O*, describing the genetic rules that govern GJ formation (Fig. 5 and Methods). We set the regularization parameter at its optimal value *α* = 0.215 (see Methods and Supporting Figure S2.). The elements of the obtained *O* matrix represent the probability that neurons expressing those genes form a GJ due to this specific genetic rule. Most of the obtained rules have a small, but non-zero weight (Supporting Figure S3), partially due to the data limitations discussed above, and partially inherent to the chosen Frobenius norm. We must therefore differentiate small values from meaningful probabilities, a challenging task because the inferred rules are not independent. To assess significance we must perform degree-preserving randomizations of the connectome [49] without violating the spatial constraints. As we lack methods to perform such randomizations without generating interactions between non-adjacent pairs of neurons, we designed a maximum entropy approach for network randomization with spatial constraints, using a subgraph randomization protocol, allowing us to determine the z-score for each predicted rule.

**FIG. 5:**
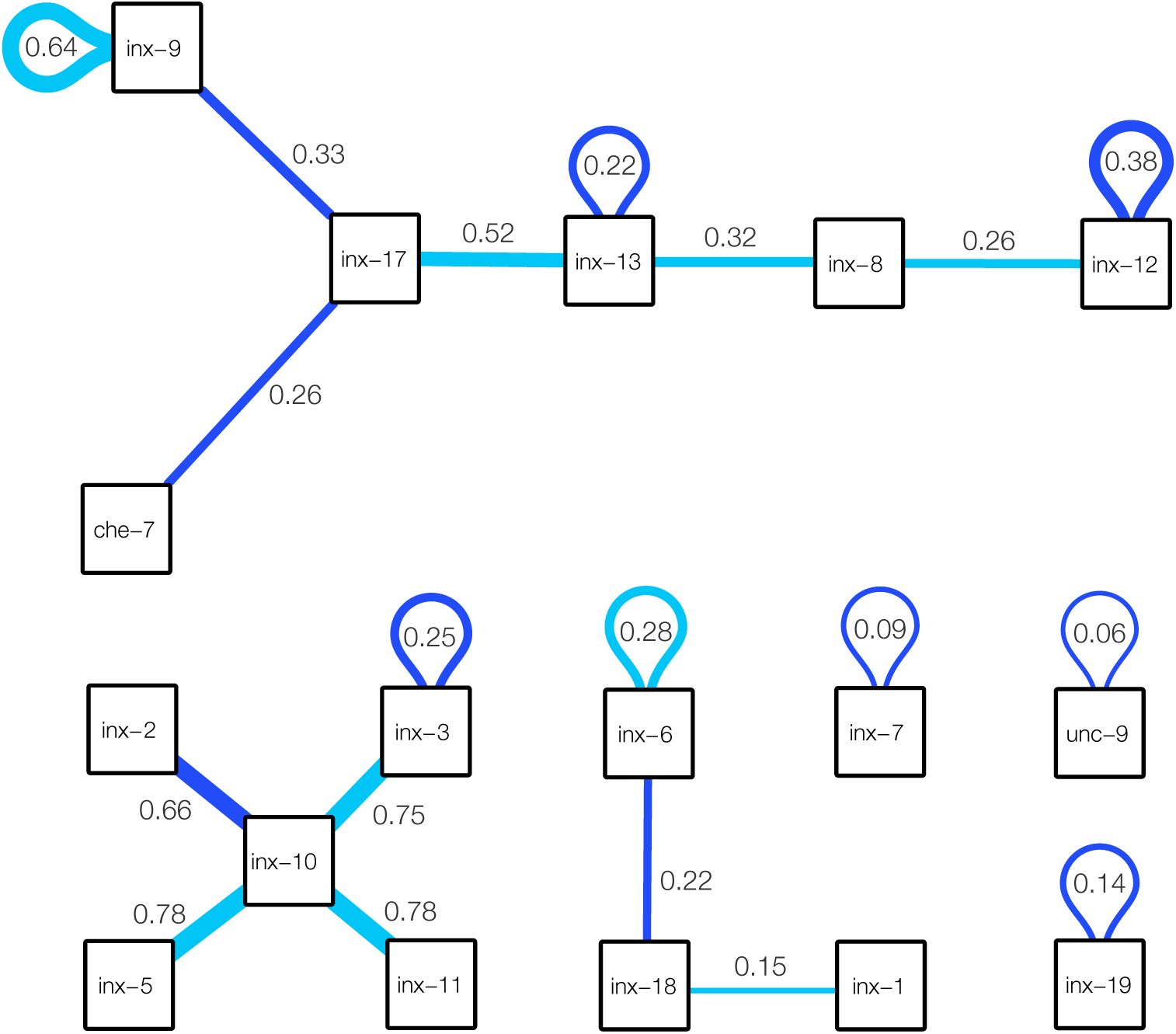
Predicted Significant Innexin Rules. Significant innexin rules inferred for *C. elegans* GJs, showing only positive rules with a z-score above 2. Each box corresponds to one of the 18 innexin proteins in *C. elegans* whose expression pattern is known. Dark blue links are found to be significant in both connectome reconstructions (Supporting Text, Supporting Figure S4), while light blue links are significant only in the Cook et al. connectome [28]. Link weights estimate the connection probability. For example the link between inx-2 and inx-10 has weight 0.66, meaning that the neurons expressing these two innexins establish GJs in 66% of the cases. Note, that the observed probability of GJs between these neurons might change if multiple rules contribute to them.

After randomization, we find 19 significant wiring rules above the threshold *z* > 2, summarized in Fig. 5. Five of the 19 rules have been uncovered previously by the experimental literature, including (1) **inx-3–inx-3** [46], (2) inx-6–inx-6 [46], (3) **inx-19–inx-19** [46], (4) **UNC-9–UNC-9** [46], (5) and **inx-10–inx-11** [50], where the boldface font indicates that the interaction is significant for two different *C. elegans* connectome reconstructions (Supporting Text, Supporting Figure S4). The inx-19–inx-19 interaction was confirmed by electrically coupling Xenopus oocytes [51], and inx-6–inx-6 channels have been confirmed by EM reconstruction [52]. The remaining interaction rules uncovered by the model are novel: (1) inx-9–inx-9, (2) **inx-9–inx-17**, (3) inx-17–inx-13, (4) **inx-13–inx-13**, (5) inx-11–inx-8, (6) inx-8–inx-12, (7) **inx-12–inx-12**, (8) **inx-2–inx-10**, (9) inx-5–inx-10, (10) inx-10–inx-3, (11) **inx-18–inx-6**, (12) **inx-7–inx-7**, and (13) **inx-17–che-7**, and (14) inx-1–inx-18, where boldface again indicates that the interaction is confirmed in both reconstructions (Supporting Text, Supporting Figure S4). Taken together, these fourteen novel interactions offer ground for direct falsifiable experimental confirmation, through genetic interventions that, according to our model, are expected to lead to rewiring in the *C. elegans* system. The developed framework allows us to predict the nature of this rewiring: for example, if a connection between two neurons is due to a single rule, then losing the participating genetic label on either side leads to a loss of interaction. For instance, our inference predicts that the AINR-ASGL and AINL-ASGR gap junctions, present in both *C. elegans* connectome reconstructions, are coded solely by inx-17–che-7 interactions, an interaction found significant according to inference on both reconstructions (Supporting Text, Supporting Figure S4). Therefore, knocking down any of these genes in the neurons, or pharmacologically preventing the interaction, is expected to result in the loss of these two gap junctions. Note that these predictions are sensitive to the noise in the input data and the choice of the significance threshold, particularly since all single-rule gap junctions originate from low-strength interactions.

## Discussion

Motivated by the need to infer the genetic rules that govern the wiring diagram of the connectome, here we have introduced a computational framework that relates the genetic expression profiles of the individual neurons to the connectome through a single brain equation (1). Although the connectome and especially the neuron gene expression profiles remain heavily incomplete and prone to noise, our results indicate that their joint coverage is sufficient to infer some of the conjectured interactions that govern gap junction formation in the *C. elegans* nervous system. To achieve this, we established a connection between the gene expression patterns of single neurons and the connectome, through the Connectome Model (1). As synapses can only form between neurons that are in physical contact, we incorporated in our framework spatial constraints, resulting in the Spatial Connectome Model (5). The model allowed us to identify 19 significant innexin rules behind heterotypic gap junctions. With the increasing availability of high quality input data, the SCM can be extended to capture heteromeric GJs and chemical synapses, illustrating the versatility of the developed modeling framework (Supporting Text).

The SCM, together with the inferred innexin rules, allows us to predict potential changes in neural wiring if gene expression is altered via knock-out experiments or silencing. Yet, a knock-out experiment of an innexin is only informative if the mutant is viable. The individual loss of several innexins (including inx-3, inx-12, inx-13, inx-14, inx-22) is known to be lethal [46], limiting knock-out experiments to non-essential innexins, unless the experiments can be limited to specific neurons only. Temperature-sensitive alleles provide an alternative way to experimentally modulate the expression of essential innexins, keeping the innexins functional during development, and disabling the corresponding gap-junctions at restrictive temperatures [53, 54]. Another possibility would be an exercise in edgetics, i.e. disrupting specific protein-protein interactions using drugs targeting innexins [55], and detecting the resulting change in the connectome. Our model could also be used to predict how the brain is rewired in the food-deprived, dormant state of the *C. elegans* known as the dauer stage. Functional studies indicate a substantial remodeling of behavior which anticipates a substantial rewiring of the GJ connectome, with profound impact on synaptic partner choices. As a prerequisite, dauer stage neuron gene expression data has been made available recently [47].

Finally, the SCM has a geometric interpretation that establishes a connection between brain connectivity and non-Euclidean geometry (Supporting Text). The rule matrix (*O*) of the SCM can always be diagonalized, leading to a set of diagonal rules in the appropriate genetic basis, given by the eigenvalues. If all eigenvalues are non-negative, that indicates that a Euclidean geometry describes the wiring rules, implying that neurons will form links to other neurons with similar expression profiles. We find, however, at least four negative eigenvalues, supporting a non-Euclidean organization (Supporting Figure S5). This indicates a strong presence of heterophily, i.e. GJ formation based on genetic complementarity rather than similarity. Such non-Euclidean wiring rules indicate that link formation in the *C. elegans* connectome is not driven by spatial proximity, rather spatiality merely constrains a fundamentally non-Euclidean structure.

## Acknowledgements

We thank Emma Towlson and Oliver Hobert for helpful discussions and Alice Grishchenko for help in designing the figures. This work was supported by the European Union’s Horizon 2020 research and innovation programme under grant agreement No. 810115 - DYNASNET. D.L.B. was supported by NIH NIGMS T32 GM008313. A.-L.B. was supported by the NSF award 1734821.

## Author contributions

I.A.K. developed the models and performed data analyses and modelling. All authors contributed to the conceptual design and the writing of the manuscript..

## Additional information

ALB is the founder of Scipher Medicine, Foodome and Nomix, that apply computational and network-based tools to health.

## Methods

### Ridge Regression

Problem (3) can be solved analytically as [44]

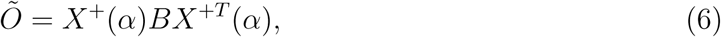

where *X*^+^(*α*) = (*X*^*T*^ *X* + *αI*)^−1^*X*^*T*^. In the *α* → 0 limit, the solution is *Õ* = *X*^+^*BX*^+*T*^, yielding the best residual error (*r*^2^) at the expense of the simplicity of *O*, prone to overfitting in the presence of errors. This limit is also sensitive to changes in *B*, therefore *α* = 0 is only appropriate when the input data is exact. In contrast, *α* → ∞ leads to the estimate *Õ* ∝ *X*^*T*^ *BX*, coinciding with the naive assumption discussed in Ref. [25], yielding a poor *r*^2^, being prone to under-fitting. Finding the optimal *α* usually requires numerical heuristics. Here we use the suggestion by Wahba [56] proven to be optimal in a generalized cross-validation scenario, corresponding to *α* that minimizes *r*^2^/*τ*^2^, where *τ* = Tr (*I* − *KK*^+^(*α*)) and *K* = *X* ⊗ *X* is calculated using the Kronecker product. Equation (5) can be solved similarly, leading to

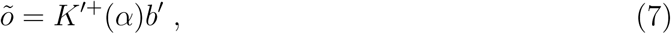

with *K*′^+^(*α*) = (K′^*T*^ K^′^+*α I*)^−1^K^*′T*^, at the optimal *α* corresponding to *τ* = Tr (*I* − *K*′*K*′^+^(*α*)) (see Supporting Figure S2).

### Subgraph Randomization

We start with a graph *G*_0_ and a subgraph G and we aim to sample uniformly the space of subgraphs of *G*_0_ with (approximately) the same subgraph degree sequence as given in G. This represents a constrained version of the traditional degree-preserved randomization, where *G*_0_ is a complete graph [57], as all interactions that are not in *G*_0_ are excluded from the randomized networks. Here, *G*_0_ represents the list of adjacent neurons that could in principle establish a GJ and we aim to randomize the network without violating the known spatial adjacency structure. We use a Maximum Entropy approach, maximizing the entropy of the random network ensemble defined as *S* = − Σ*P*(*G*) ln *P*(*G*), where the average degree of each node is then ⟨*k*_*i*_⟩ =Σ_*G*_ *P*(*G*)*k*_*i*_(*G*), which we aim to keep fixed to their original value, *k*_*i*_. The probability of a given graph instance is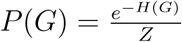, where *H* = Σ_*i*_ *β*_*i*_*k*_*i*_*(G)* and the probability of having a link between nodes *i* and *j* can be expressed as 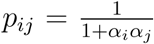 where 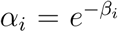. The average degree of a node is then given by 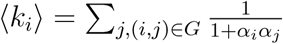, and the optimal *α* can be found iteratively, with the update rule

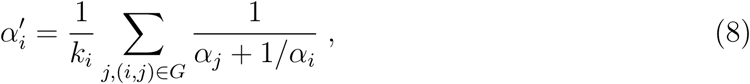

starting from the initial condition 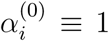 leading to 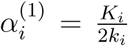. We perform at least a hundred iterations to estimate the optimal *α*, allowing us to calculate the mean and the standard deviation of the randomized matrix ensemble. Due to the linearity of the SCM solution, these yield a z-score for each inferred wiring rule.

## Supplementary Text

### Available Connectomes

The neural system of *C. elegans* consists of chemical synapses and gap junctions between 302 neurons. Graph theoretical analyses of *C. elegans* connectomes typically restrict the circuit to the connected somatic nervous system of 279 neurons, excluding 20 neurons in pharyngeal nervous system and 3 somatic neurons (CANL/R and VC06) that, in the Varshney et al. reconstruction [1], do not synapse with other neurons. We studied two different reconstructions:

### Cook et al

reconstructs a gap junction network with 1,051 undirected links, including 11 self-loops [2].

### Varshney et al

provides a more conservative reconstruction, resulting in a gap junction network has 517 links, of which 3 are self-loops [1].

Given its updated methodology, the Cook et. al. connectome is utilized in the body of the paper. However, we also performed the SCM analysis on the Varshney connectome (Supporting Figure S4), which resulted in 11 interactions that were shared with the Cook et al. analysis: (1) inx-3–inx-3, (2) inx-9–inx-17, (3) inx-19–inx-19, (4) UNC-9–UNC-9, (5) inx-10–inx-11, (6) inx-13–inx-13, (7) inx-12–inx-12, (8) inx-2–inx-10, (9) inx-18–inx-6, (10) inx-7–inx-7, and (11) inx-17–che-7.

### Heteromeric Gap Junctions

As documented in vertebrate connexins, two or more innexins might form mixed, heteromeric hemi-channels, playing an important role in building the connectome. Therefore, innexins are also expected to form heteromeric GJs [3], meaning that in a given neuron the expression of two (or more) innexins are required to establish a GJ with another neuron [4]. Heteromeric hemi-channels formed by two kinds of innexin proteins can be incorporated in our model by adding the joint (logical AND) expression of innexin pairs to *X* as additional labels. However, even at the pairwise level, such a step increases the dimensionality of the problem by an order of magnitude (from 18 to 171 labels) for *X* and from 171 potential rules to 14, 706 rules in *O*. This implies that we have far more rules to extract than the number of mapped GJ connections, rendering the problem seriously underdetermined. To avoid overfitting, a potential strategy is to add each innexin pair individually to the innexin labels to see which pair improves the most on the model’s residual error. To avoid false positives, the inference of heteromeric rules is left as future work, gaining relevance once the homomeric rules have been experimentally validated.

### Chemical Synapses

Given that the formation of GJs is governed by innexin expression, we know the precise genetic basis in which the rules operate. To perform the same analyses for chemical synapses, we must extend the search basis to the entire list of genes, as any gene could contribute to the identity of the neurons. In other words, unless the correct biological basis can be further restricted, we lack the labels corresponding to the combination of proteins contributing to synapse formation. One way to reduce the dimensionality of the problem is to focus on transcription factor (TF) expression only. However, as opposed to innexin expression, adult TF expression might not be strongly related to the expression history during development, shaping the chemical synapses. Ideally, we would need to restrict the number of labels to at least the number of neurons, to infer the chemical rules using the procedure we followed for innexins. However, even then, given that synapses are directed, we need two sets of labels, one for identifying the input neurons (*X*) and one for the output neurons (*Y*), extending the Connectome Model into a Directed Connectome Model (DCM)

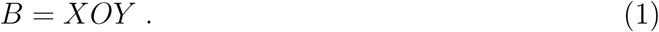

The DCM can be solved similarly to the CM as *Õ* = *X*^+^*BY*^+^.

### Geometric Interpretation of the Wiring Rules

Predicting the wiring rules (*O*) from *B* and *X* (Problem II) represents an inverse geometric embedding problem. Indeed, we have N input vectors X_*i*_, one for each neuron, where each vector represents the expression profile of M genes within that neuron. Hence, the expression pattern of each neuron can be interpreted as a position in an M-dimensional space. In a three dimensional Euclidean space, the scalar product ⟨*x, y*⟩ = *x*_1_*y*_1_ + *x*_2_*y*_2_ + *x*_3_*y*_3_ measures the overlap of the two vectors x and y. Hence ⟨*x, y*⟩ = 0 for vectors that are orthogonal to each other, and ⟨*x, x*⟩ gives the squared length of the vector x. We can formally generalize the scalar product as an abstract pseudo-metric, defining the overlap between two vectors as 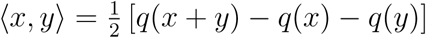, reducing to the Euclidean metric for 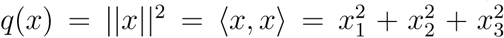. In the light of this geometric interpretation, the r.h.s of Eq. (1) defines a scalar product 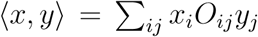, where the choice of *O* guarantees that there is a non-zero overlap between neurons pairs, x and y, that satisfy the genetic condition for synapse formation. In this picture, non-interacting neurons correspond to orthogonal vectors, with ⟨*x, y*⟩ = 0, according to the pseudo-metric defined by the wiring rules in O. The Euclidean metric would imply that neurons form synapses whenever they share at least one expressed gene (or share a genetic label). In this case *O* would be a diagonal matrix, and it would be unable to describe the combinatorial rules described in Figs 1 and 2 required for synapse formation. Hence, instead of a diagonal matrix, we need to use an *O* capable of capturing the combinations of genes needed for synapse formation, like the one shown in Fig. 2b. For an Euclidean metric, all eigenvalues are positive (or zero). The existence of at least one negative eigenvalue indicates that the system is embedded in a Minkowski-like space, encountered in special relativity, where 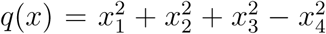, with eigenvalues (1, 1, 1, −1), the fourth coordinate representing time *x*_4_ = *ct*.

As (5) allows us to infer the genetic rules governing GJ formation through the expression of 18 innexin proteins, we are able to unveil the 18 dimensional geometry behind GJs. We find that in *C. elegans*, the genetic rule matrix, *O*, has as at least four negative eigenvalues (Supporting Figure S4), indicating a Minkowski-like space with at least four time-like dimensions. In Euclidean space the triangle inequality is fulfilled (||*x*+*y*|| ≤ ||*x*||+||*y*||), while in Minkowski-like spaces it is violated in some directions, leading amongst others to the *twin paradox*. The triangle inequality appears as the popular *triadic closure principle* (TCP) of social networks, where two friends of a node are also friends with each other, with high probability. In a non-Euclidian space, the friend of my friend is not necessarily my friend anymore. The negative eigenvalues of *O* indicate that the triangle closure principle is violated, hence similarity and complementarity together drive GJ formation. Let’s illustrate this for a single genetic rule. The inx-2–inx-2 self-interaction would be consistent with a Euclidean structure, where GJs are formed when two neurons both express inx-2, representing expression profile similarity. In contrast, the wiring rule inx-2–inx-3 can be only interpreted in a Minkowski space, where similar neurons do not necessarily interact, but we need complementary expression, i.e. the expression of inx-2 in one neuron and inx-3 in the other, for GJ formation. In general, both similarity and complementarity is expected to play a role in neuronal wiring. Note, that we could not have arrived to this conclusion by inspecting the spectrum of the connectome B, given that *B* is truncated by spatial constraints.

To conclude, we find that the genetic rules of the connectome span a high dimensional pseudo-Euclidean space, despite the embedding of the physical connectome in a three-dimensional Euclidean space.

---

**FIG. S1:**
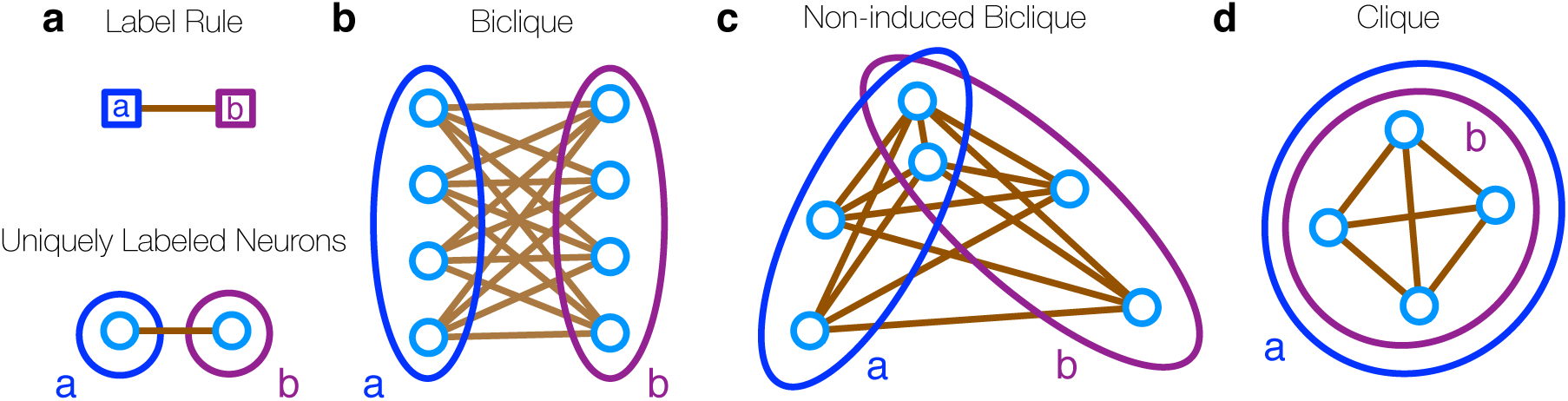
Topologies resulting from a single, deterministic rule. A genetic rule, connecting two labels, can appear very differently in the connectome depending on the distribution of the labels in the network. Each rule *O*_*ab*_ linking two labels (*a* and *b*) encodes *E*_*ab*_ = *O*_*ab*_ (*N*_*a*_*N*_*b*_ − *N*_*ab*_(*N*_*ab*_ − 1)/2) neural connections, where *N*_*a*_ and *N*_*b*_ are the number of neurons with labels *a* and *b*, and *N*_*ab*_ is the number of neurons expressing both labels. **a)** *A* rule can correspond to a single neural connection if the labels identify individual neurons. **b)** If the labels extend over multiple neurons but share no common neurons, we have a biclique in the graph. **c)** If the labels overlap for some neurons, we observe a generalized, non-induced biclique, reminiscent of core-periphery structure. **d)** If the labels overlap completely, a clique is formed.

**FIG. S2:**
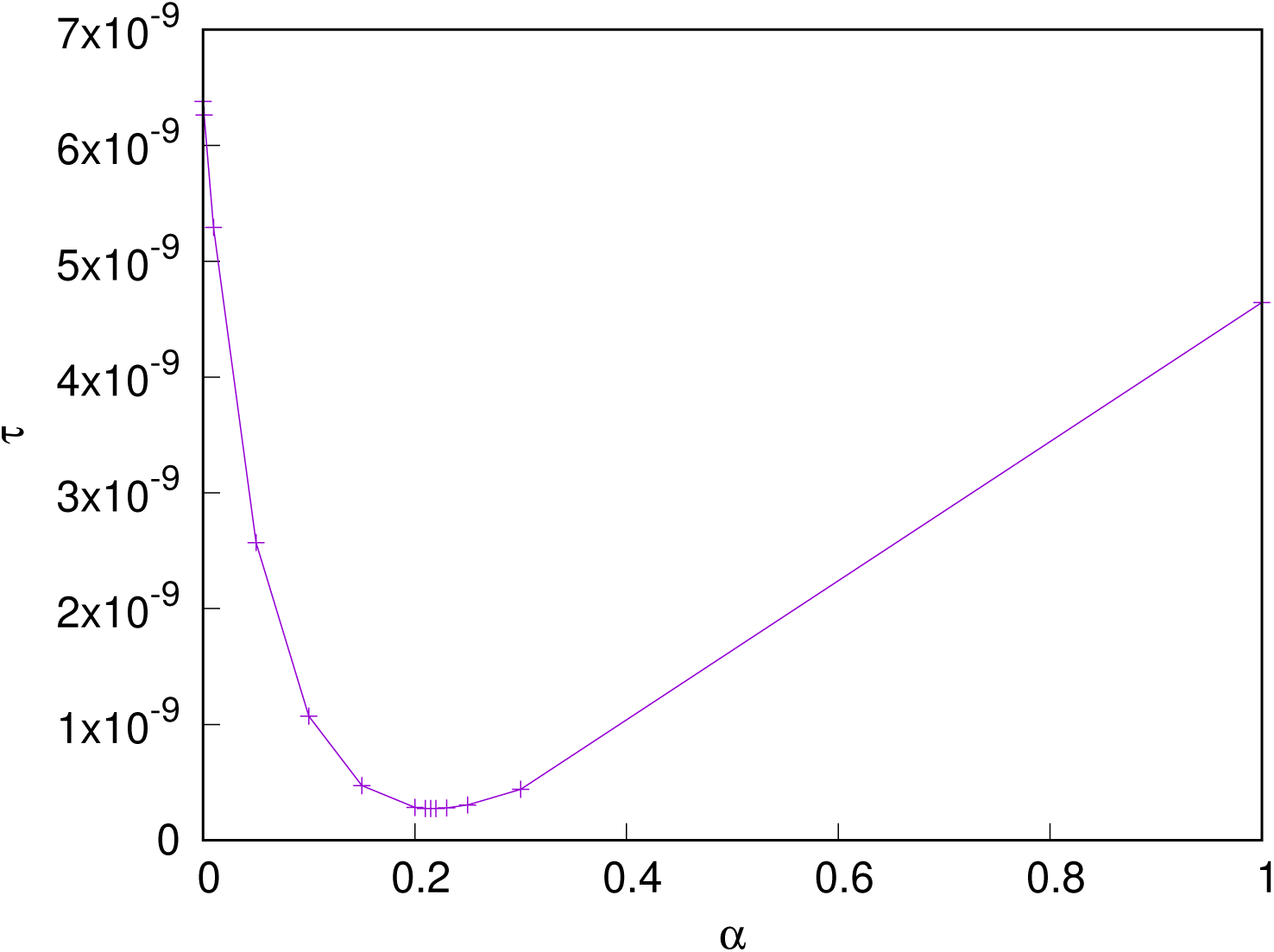
The optimal *α* = 0.215 regularization parameter for the SCM. The *y*-axis shows the *τ* value in addition to a constant background of *τ*_0_ = 8.55 × 10^−6^.

**FIG. S3:**
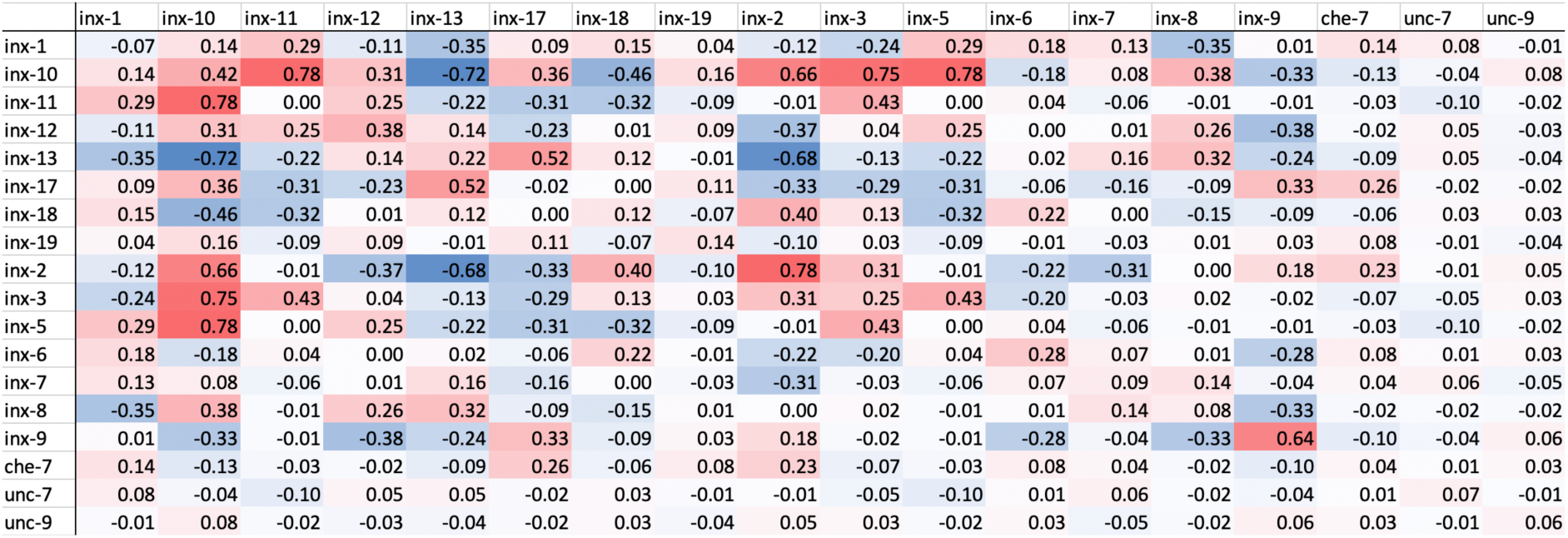
Solution of the SCM for Cook et al. data. Positive rules are indicated in red while negative rules in blue.

**FIG. S4:**
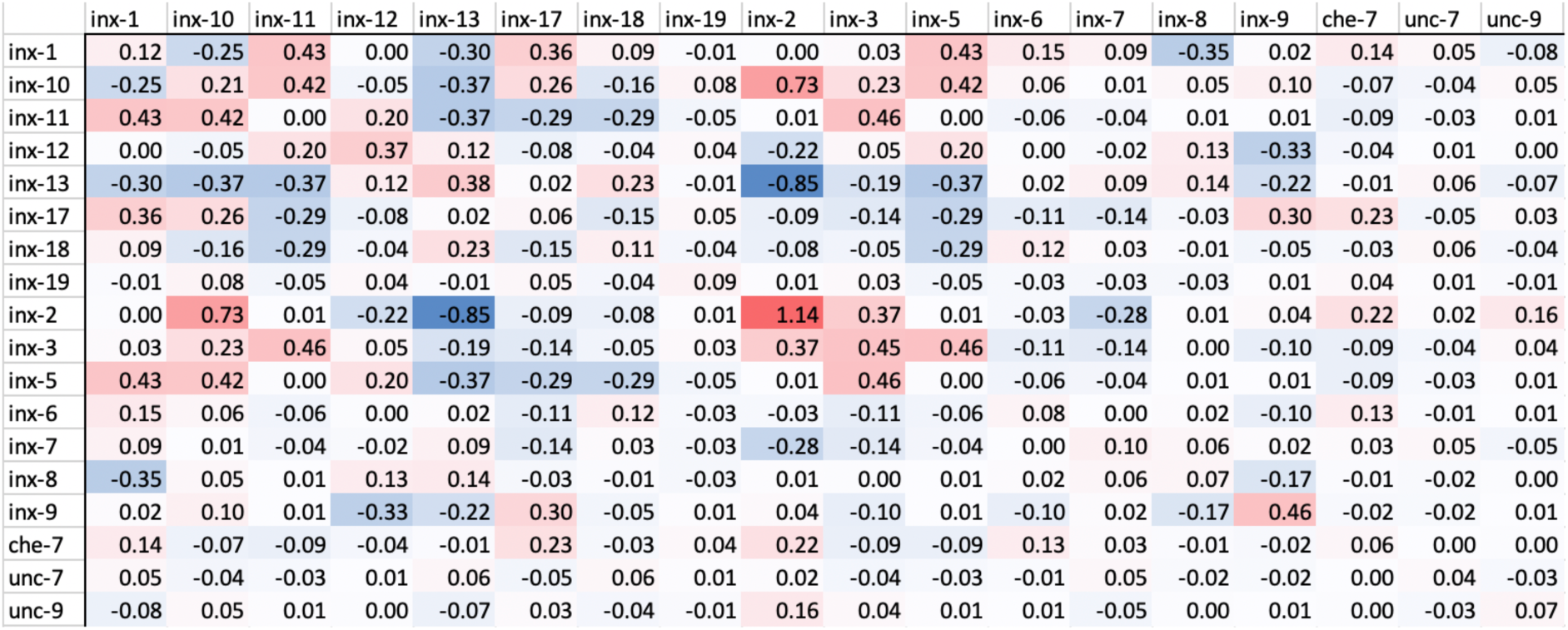
Solution of the SCM for Varshney et al. data. Positive rules are indicated in red while negative rules in blue at the optimal *α* = 0.12 regularization parameter value.

**FIG. S5:**
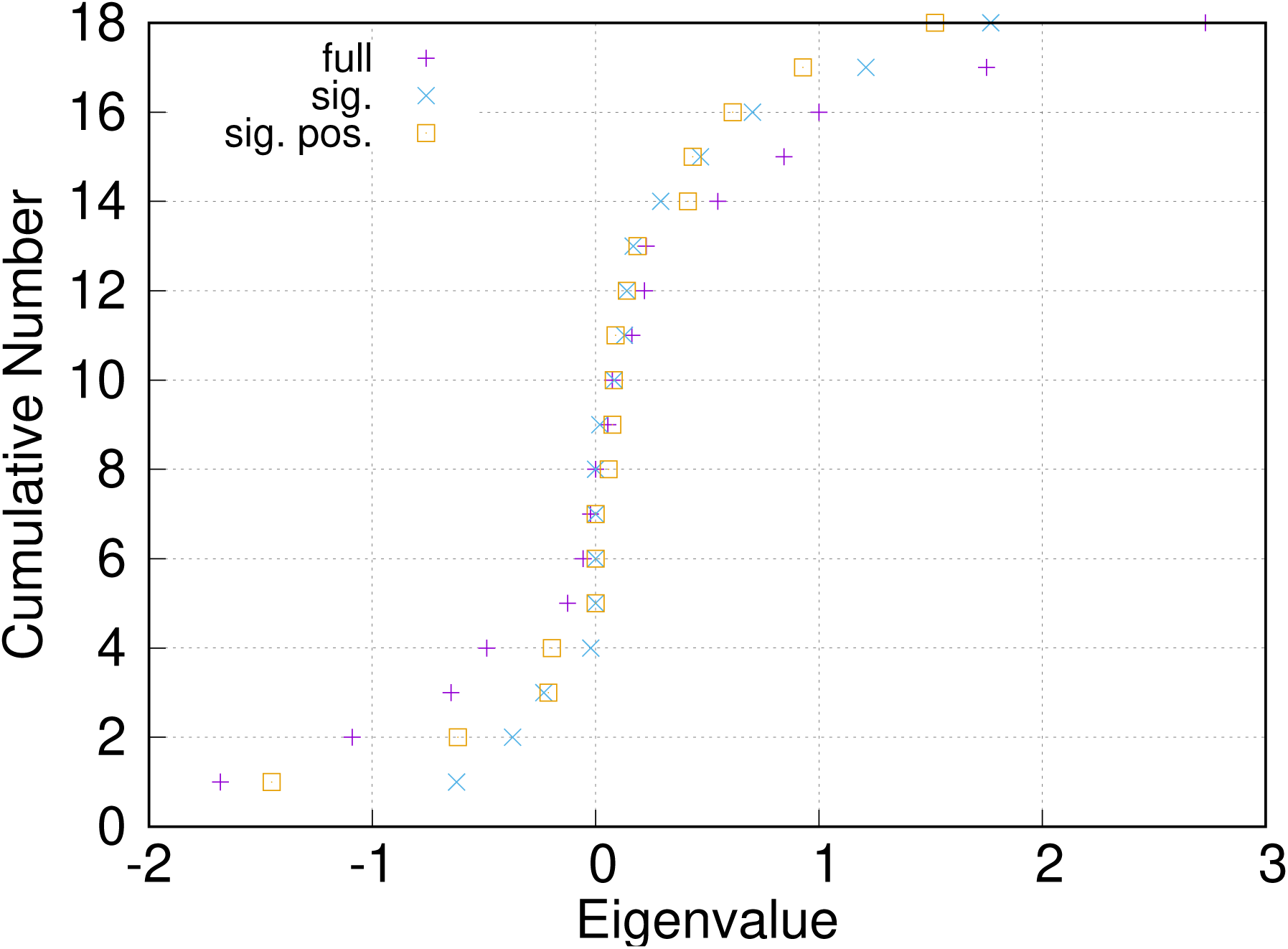
Eigenvalues of *O*. The obtained genetic rules have an indefinite spectrum as shown for the obtained *O* matrix, keeping all entries (full), only the significant ones with |*z*| > 2 (sig.) or only the significant positive entries (sig. pos.).

